# Human cerebral tissues created via active cellular reaggregation produce functionally interconnected 3D neuronal network to mimic pathological circuit disturbance

**DOI:** 10.1101/2020.06.11.144832

**Authors:** Aref Saberi, Albert P. Aldenkamp, Nicholas A. Kurniawan, Carlijn V.C. Bouten

**Author notes:** **Correspondence and requests for materials** should be addressed to N.A.K.

## Abstract

Various characteristics of a developing central nervous system, including sensory input-output^1,2^, neuronal migration^3,4^, and regionalization^3,5^, have been recapitulated through recent advances in the culture of brain organoids. These organoids can model the fundamental processes in brain development and disease. A remaining critical challenge, however, is to achieve complex neuronal networks with functional interconnectivity as in native brain tissue. Generation of current organoid models originates from classic dissociation-reaggregation paradigms^6^, often relying on mechanically-enforced quick reaggregation of pluripotent stem cells^7^. Here we describe an alternative method that promotes matrix-supported active (migrative) reaggregation of cells (MARC), reminiscent of in vivo developmental morphing processes, to engineer multi-regional brain tissues in vitro. Measurements of neuronal activity in intact 3D tissues revealed functional interconnectivity, characteristic of cerebral neuronal networks. As a proof of concept, we show that interconnected cerebral tissues produced using this approach can mimic propagation of epileptiform discharges in a custom-built in-vitro platform.

## Main body

From a developmental perspective, one key step in the assembly of neuronal circuits and mature interconnected networks is neuronal migration^8^. This step is, however, inherently suppressed in the often used protocols to generate brain organoids (e.g., based on serum-free culture of embryoid body-like aggregates with quick reaggregation or SFEBq method^7^). In order to recapitulate this process, we introduce a novel culturing method to develop human cerebral tissue at the mesoscale (mm size) via matrix-supported active (migrative) reaggregation of cells (MARC). Unlike other protocols, e.g. reported for brain organoids, that use a quick, mechanically-enforced aggregation of the dissociated cells, the formation of the three-dimensional (3D) tissue (hereinafter referred to as “cerebral tissue”) is initiated by active (migrative) reaggregation of the cells during chemically-induced differentiation, with the immediate 3D extracellular support of Matrigel (Figure 1a; see also experimental details in Methods). In order to promote active neurite-mediated reaggregation and engineer multiregional cerebral tissues, we rationally designed a neuronal differentiation protocol employing phased introduction and withdrawal of the culture additives used for neuronal tissue patterning. After ~80% confluence of the culture of human induced pluripotent stem cells (hiPSCs) under feeder-free condition, neuronal differentiation was initiated by addition of SMAD inhibitors (dorsomorphin and SB431542), a GSK-3 inhibitor (CHIR99021), SHH, and b-FGF. After 7 days of induction, the cells were enzymatically dissociated and homogenously resuspended in a mixture of Matrigel and neural differentiation medium containing b-FGF, SHH and FGF8. This step is henceforth considered Day 0 of MARC culture. Neural differentiation of 7 days therefore took place in a 3D environment, accompanied by the rapid formation of spheroids with a size of 200–300 μm (Fig. 1b, Day 7). Pre-terminal differentiation was started by removal of b-FGF from the medium, resulting in the formation of neurite outgrowth and bundles that connected the spheroids to each other (Fig. 1b, arrows and arrowheads). These spheroids merged over time (approximately 2 weeks), likely through a synapse-mediated migration ^9^, resulting in large cerebral tissues with a size of 2–4 millimeters (Fig. 1b, Day 15, 20). Finally, SHH and FGF8 were withdrawn and the cultures were treated with the maintenance medium. The cerebral tissues continued growing during the culture period of 90 days and expressed markers of distinct cell types of the human brain, confirming multiregional tissue patterning of our engineered cerebral tissues (Figure 1c and Figure S1). In addition, colocalized expression of different markers suggested spatial patterning of these cell populations (Figure S1).

**Figure 1.**
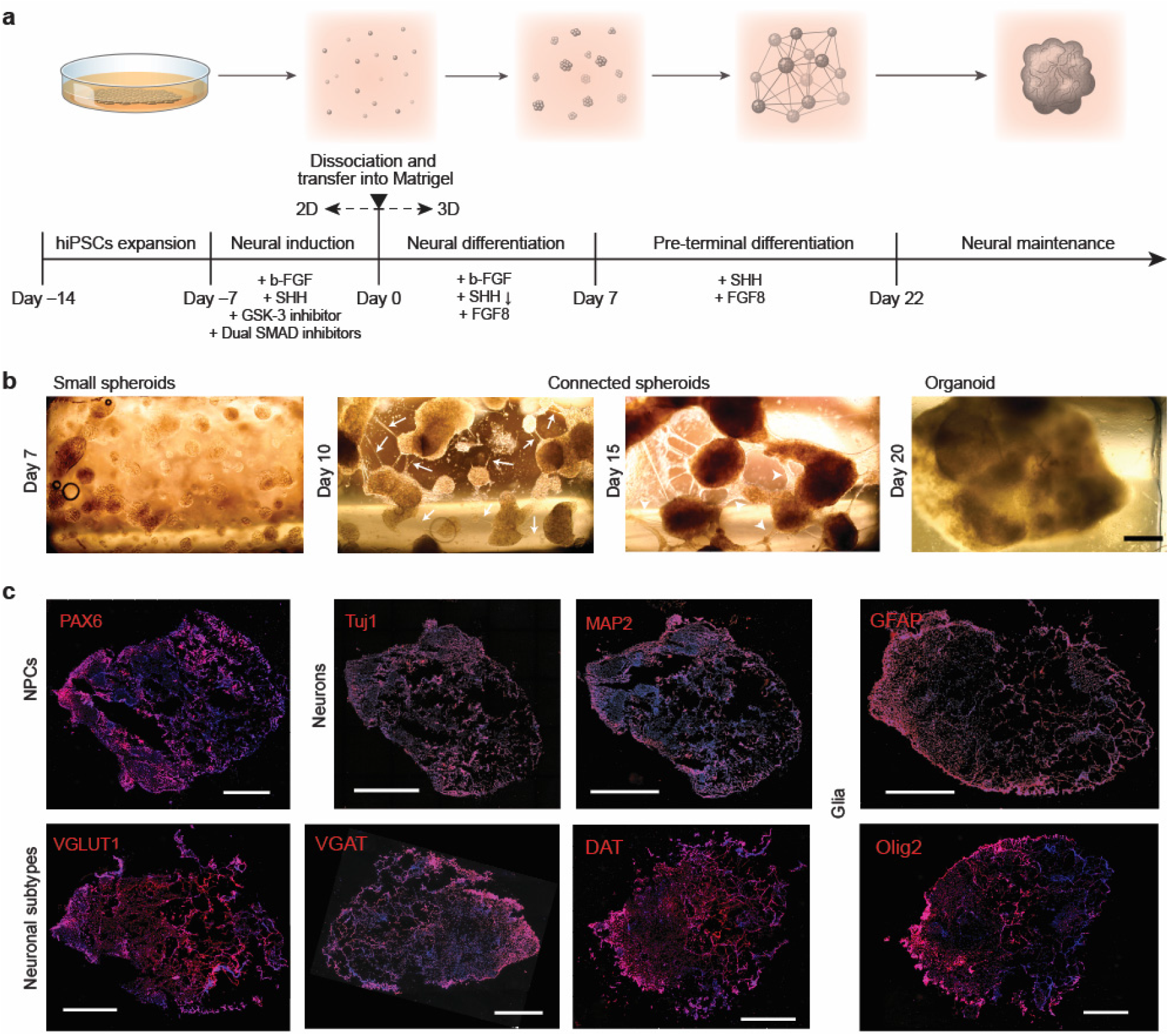
Formation of human cerebral tissues via MARC. (**a**) Schematics of the MARC culture method, showing the different culture steps. Timeline and additives supplied in each step are indicated. (**b**) Example phase-contrast images at the different 3D-culture phases, showing the matrix-supprted active reaggregation of the cells into cerebral tissues. Single dissociated cells suspended in Matrigel grew into small spheroids (Day 1–7). During pre-terminal differentiation, neurite outgrowths extended from the spheroids (white arrows) and merged into neurite bundles (white arrowheads) between spheroids (Day 10, 15). The spheroids migrated using these neurite bundles and merged into large cerebral tissues (Day 20). Scale bar: 500 μm. (**c**) Immunohistochemical staining of cerebral tissues at Day 90 revealed the presence markers of neural progenitor cells (NPCs; PAX6), early and mature neurons (Tuj1 and MAP2), mature GABAergic neurons (excitatory VGLUT1 and inhibitory VGAT transporters), mature dopaminergic neurons (DAT), astrocytes (GFAP), and oligodendrocytes (Olig2) were expressed (red), indicating multiregional cerebral tissues with extensive cellular diversity. Blue = DAPI. Scale bars: 1 mm.

To test the neuronal functionality of MARC-produced cerebral tissues, we evaluated the neuronal interconnectivity within the intact 3D tissues using live calcium imaging. We observed that cerebral tissues at age 4 weeks exhibited extensive spontaneous calcium surges throughout the tissues (Fig. 2a,b and Supplementary Movie 1). Examination of the time-lapse images also indicated extensive synchronized neuronal firing, which is known to result from concentrated bursts of action potentials between interconnected neurons, leading to influx of extracellular calcium^10^. To quantify this population-wide intercellular synchronized activity, we computed the pairwise linear correlation coefficient *r* from the intensity time-trace of 387 regions-of-interest (ROIs) representing detected single neurons in the cerebral tissues. The correlation matrix between all ROI pairs shows that the majority of the ROI pairs had low *r*-values, but a significant number of pairs was highly correlated (Fig. 2c). We defined two ROIs to be functionally connected when *r* > 0.6, following Eguiluz et al^11^. Functional neuronal connection was found for 304 pairs (~0.4% of all ROI pairs analyzed) and depicted in a spatial connection map (Fig. 2d), demonstrating a functional neuronal network. Interestingly, the functional connections were not randomly distributed throughout the cerebral tissues; several ROIs were highly connected to many other ROIs whereas most others only had a few connections. In fact, we found an increased synchrony between the nodes with higher amount of connections (Fig. 2e). These observations are consistent with the topological features of scale-free networks, which have been proposed to be important for synchronized functional networks^12–14^. To check whether the functional interconnectivity in the cerebral tissues follows a scale-free topology, we plotted the distribution of the number of connections each neuron has. Indeed, the distribution follows a power law with a decay constant of −2 (Fig. 2f), consistent with the characteristics of scale-free network^15^ and in agreement with clinical measurement of whole-brain activity^16^. The presence of a small number of hyper-connected “hub-like” cells (up to >20 connections in our case) and a large number of cells with few connections results in a low average number of connections (~1.5 in our cerebral tissues). It has been proposed that brain achieves large-scale interconnectivity between brain regions despite the low average number of connection through a modular network topology, whereby the network is composed of subnetworks (“modules”) of densely interconnected neurons (“nodes”). Such modular architecture is thought to be critical for the emergence of adaptive behaviors and cognition^17,18^. To assess the network modularity in our cerebral tissues, we analyzed the functional connectivity using iterative Louvain community-detection algorithm^19,20^ (see Methods). The algorithm identified 3 distinct modules within the tissue with sparse intermodular node connections (Fig. 2g,h). Moreover, each module includes its own local hubs that are highly interconnected (inset in Fig. 2h), which ensure global integration of functional interconnections across the overall network^21^. Within each module, the nodes are interconnected in a hierarchical topology, from a few hub nodes with high number of connections that are closely connected to each other to peripheral nodes at the outer edges of the network topology (Fig. 2i). Taken together, the analysis demonstrates a rich interconnectivity in the cerebral tissues reminiscent of the emerging attributes of functional networks in brain, suggesting their utility as an in vitro mimic of brain functional interconnectivity.

**Figure 2.**
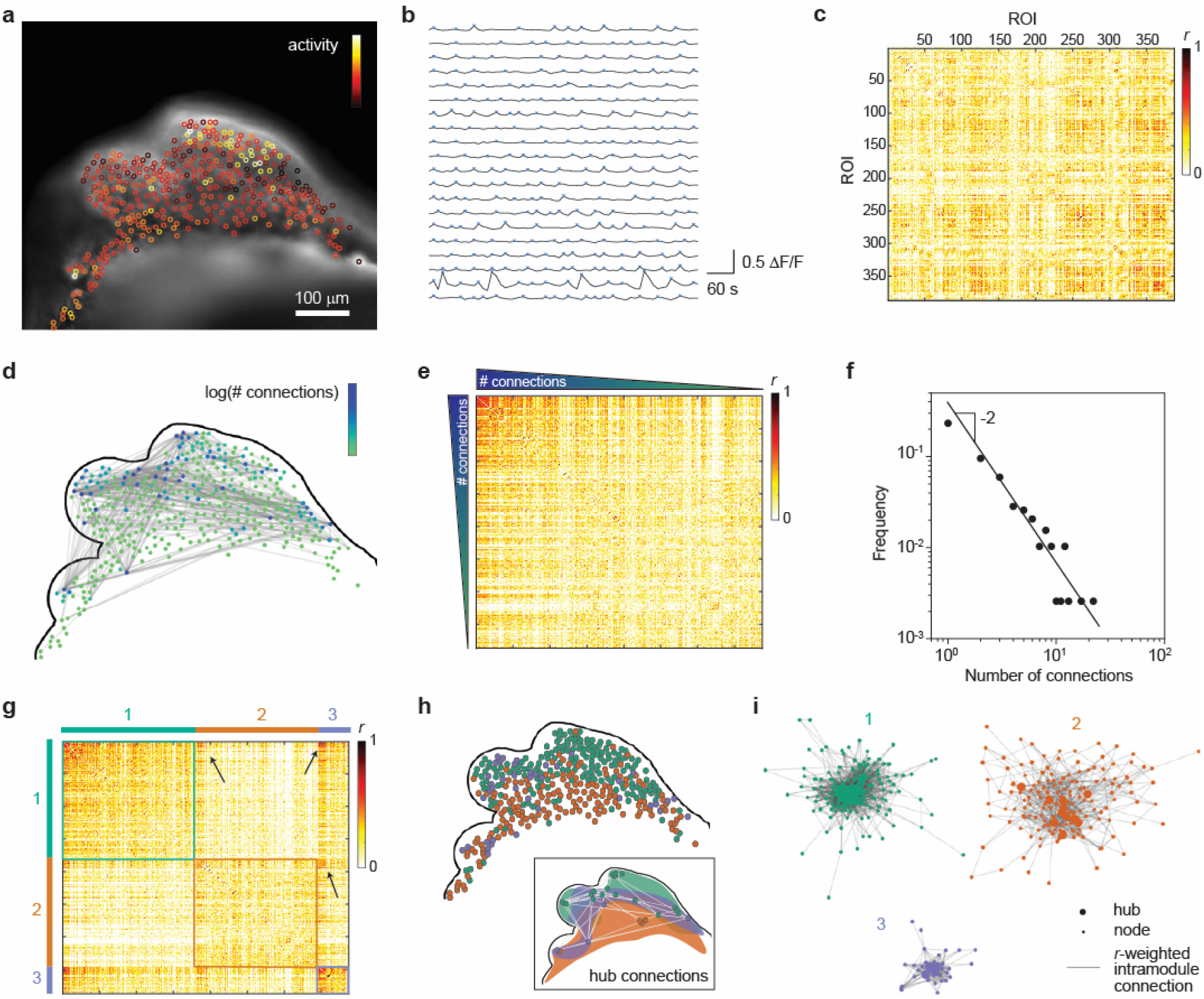
Functional network interconnectivity in intact MARC-produced cerebral tissues. (**a**) Neuronal activity in a cerebral tissue at week 4 of MARC culture. A snapshot of live fluorescence calcium imaging on the intact cerebral tissue is overlaid with regions-of-interest (ROIs) color-coded based on the frequency of detected calcium surges (“activity”) in each ROI. See Supplementary Movie 1 for the corresponding time-lapse imaging data. Scale bar: 100 μm. (**b**) Representative time traces of normalized intensity (ΔF/F) from 20 ROIs. Blue crosses indicate detected transient spikes. Time traces from 387 ROIs in the image are used to compute the correlation coefficient *r*_ij_ between ROI pairs *i* and *j* (see Methods for details of calculation). (**c**) Correlation matrix between any pair of the 387 ROIs, showing the *r*-value for each pair. (**d**) Depiction of functional connectivity network in the cerebral tissue. The contour of the cerebral tissue, corresponding to **a**, is shown. Two ROIs are defined to be functionally connected when *r* > 0.6 and shown as gray lines in the connectivity map. Further, each ROI in the connectivity map is color coded based on the number of functional connections it has. (**e**) The same correlation matrix as in **c**, but with the ROIs sorted based on the number of functional connections they have. The high density of ROI pairs with high number of connections and high *r*-value suggests a non-random network topology. (**f**) Distribution of the number of functional connections. The distribution follows a power law with a decay power of −2, demonstrating a scale-free cerebral tissue functional network. (**g**) Clustered correlation matrix. To test whether the network exhibits modular topology, the functional connectivities are analyzed using Louvain algorithm^19^, which indicates that the network contains three communities/modules. The correlation matrix in **d** is then reordered so that the nodes in the same module (color coded with 1, 2, and 3) is positioned together. The high density of cross-module node pairs with high *r*-values (arrows) suggests the existence of hub connections between modules. (**h**) The localization of the nodes in the 3 modules. The color of the nodes correspond to the color coding of the 3 modules in **g**. The inset shows the spatial regions that enclose the nodes identified in the 3 modules, together with the hub nodes (defined as nodes with number of intra-module connections larger than 90^th^ percentile in the module) and the cross-module hub connections (gray lines). (**i**) Topological representation of the intra-module functional connectivity networks. To illustrate the topological proximity of highly connected nodes, each module network is shown using Fruchterman–Reingold algorithm^22^, where the length of the lines connecting nodes is proportional to 1–*r* (i.e. short lines indicate high correlation coefficient between the node pairs, and the converse). The hubs in each module are also indicated. The central positioning of the hub nodes, as well as the close topological proximity between the hub nodes, highlight their status as intramodular connector nodes in the cerebral tissue functional network.

Having established the functional interconnectivity within the MARC-produced cerebral tissues, we wanted to study 3D interconnectivity and signal transmission between living, interconnected cerebral tissues. To this end, we designed and fabricated a polydimethylsiloxane (PDMS) chip with optimized features, consisting of two culture chambers that are individually accessible and separated by a porous membrane (Fig. 3a). The design and dimensions of the vertically-tapered chambers were chosen to suit one-pot formation of MARC-produced cerebral tissues and to maintain nutrition and oxygen supply, while at the same time allowing simultaneous visualization of both chambers in a side-by-side configuration through a glass slide at the bottom (Fig. 3b). The porous membrane separating the chambers,was chosen to have pore sizes of 8 μm to keep cerebral tissues separated yet allow spontaneous neurite interconnection across the membrane. Following the MARC culture protocol, we generated cerebral tissues in each of the two chambers, which formed in a similar way as described earlier, including the neurite-assisted spheroid reaggregation into cerebral tissues (Fig. 3c). Importantly, the tissues were observed to spontaneously connect with each other through the porous membranes (Fig. 3d). Live calcium imaging showed frequent calcium surges across the membrane (Fig. 3e and Supplementary Movie 2), indicating the presence of functional interconnectivity between the tissues in the two chambers. Therefore, the combination of cerebral tissue formation using MARC and this interacting separated 3D coculture (iS3CC) chip is ideally suited for investigations into the signal transmission between interconnected cerebral tissue cultures.

**Figure 3.**
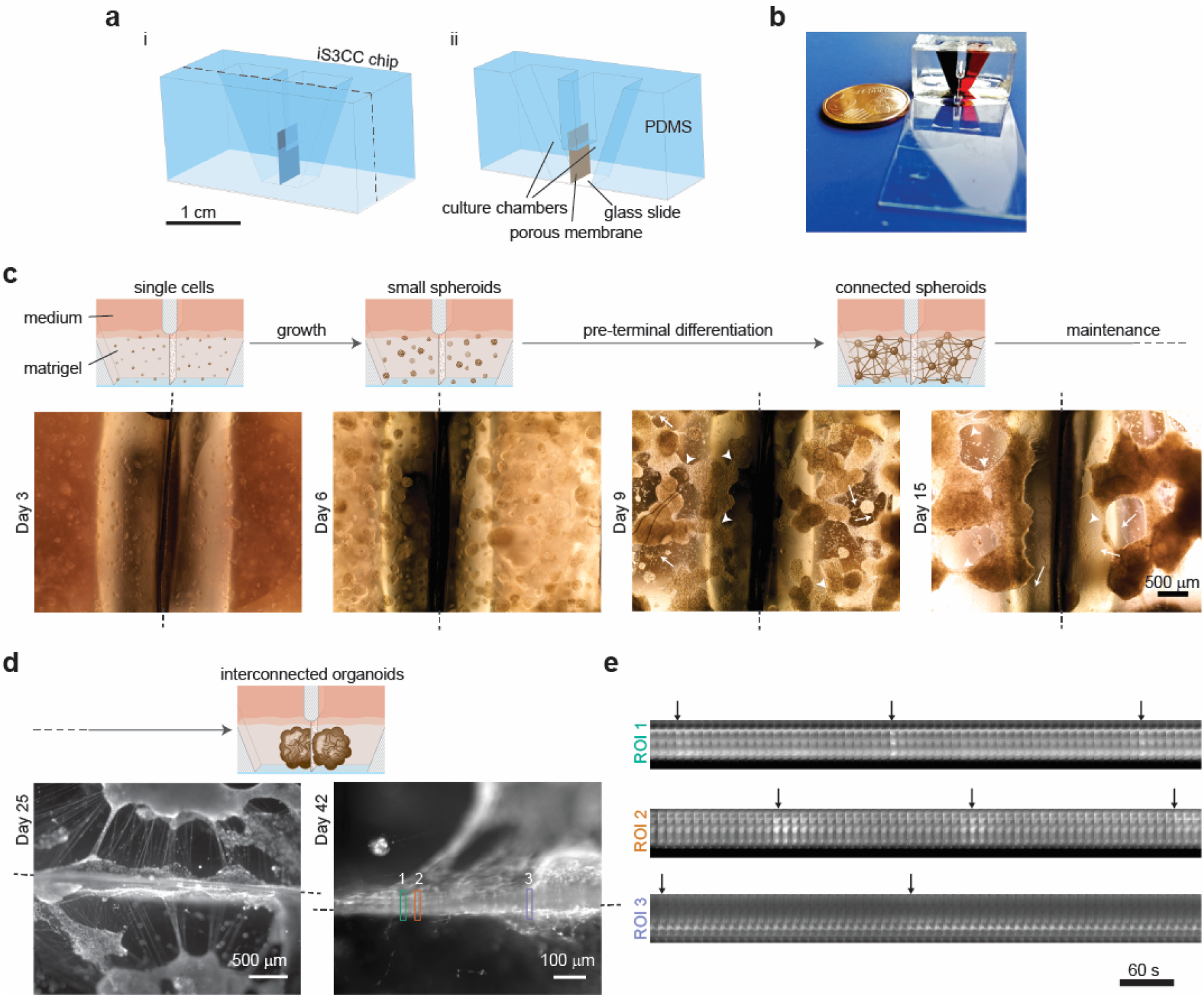
Formation and interconnection of MARC-produced cerebral tissues in the iS3CC chip. (**a**) The design and features of the iS3CC chip. A schematic illustration of the iS3CC chip (**i**) and a cross-section view of the iS3CC chip including the features: the PDMS body, chambers wherein the cerebral tissues are cultured, the porous membrane, and the glass slide (**ii**). (**b**) A photograph of an assembled iS3CC device where chambers are filled with red and blue dyes. (**c**) Side-view schematics of the progress of MARC culture in the iS3CC chip, resulting in interconnected cerebral tissues. Bottom-view phase-contrast images of both chambers of the iS3CC chip shown at the bottom, demonstrating daily progress of MARC culture during different phases of cerebral tissue formation. White arrows indicate connective neurite outgrowths, whereas white arrowheads indicate merged neurite bundles between spheroids. Dashed lines indicate the porous membrane separating the chambers. (**d**) Fluorescence pictures of intracellular calcium detected by fluo-4 direct, in cerebral tissues at day 25 and 42 and extended neurite outgrowths and bundles from both separated cultures, across the porous membrane indicated by horizontal dashed lines. (**e**) Live calcium imaging in both interconnected cerebral tissues in the chambers of the iS3CC chip, demonstrating active connections between the separated cerebral tissues. The kymographs of three ROIs (shown in corresponding color code in **d**, right) show neural activity of connections across the membrane as a function of time. Black arrows indicate the calcium transients during the imaging time. Scale bar: 60 s.

Finally, as a proof of concept of our approach, we sought to demonstrate its utility to mimic a neurological disorder affecting network interconnectivity in a controlled *in vitro* environment. Epilepsy is a chronic network-level disease defined and diagnosed by the occurrence of one or more unprovoked epileptic seizures, which are caused by alterations in the brain network circuits and functional interconnectivity. Physiologically, epileptic seizures are characterized by a transient occurrence of abnormal excessive or synchronous neuronal activity and spatial propagation of these abnormal activities. To experimentally induce epileptic seizures, we used the neurotoxic properties of Penicillin G^23^, a γ-aminobutyric acid (GABA) A-receptor (GABA_A_R) blocker of the β-lactam antibiotics family^24^. The epileptiform mechanism of Penicillin G has been theoretically and experimentally shown to occur through the specific binding of Penicillin G with GABA_A_R in an open configuration, which prevents GABAergic transmission in CNS^25^ and results in a hypersynchronous activity in the brain due to interference of the GABA-inhibition and glutamate-excitation equilibrium, causing abnormal electrical discharges^26^. Here we intended to simulate propagation of abnormal discharges from one cerebral tissue to another. Therefore, we formed interconnected cerebral tissues in the two chambers of the iS3CC chip, treated one chamber with Penicillin G by bath application to the culture medium (Fig. 4a), and monitored the neuronal activity in both chambers using live calcium imaging (Fig. 4b,c). Prior to the Penicillin G treatment, both cerebral tissues showed comparable level of neuronal activities (Fig. 4b,c inset). Immediately following the Penicillin G treatment, we observed an increased amount of fluorescence in the treated tissue, compared to the baseline (Figure 4b,g, Figure S2, and Supplementary Movie 3). This is accompanied by an increased magnitude and frequency of neuronal activity, as well as synchrony of transient spikes in the treated tissue (Figure 4e,f,h, in blue), indicative for abnormal excessive discharges. Subsequently, the neurons in the untreated chamber similarly showed an increased intensity and neuronal activity post-treatment (Fig. 4c,g), despite negligible inter-chamber particle diffusion (Figure S3). This observation strongly suggests that these immediate changes in the untreated tissue result from discharge propagation through intertissue connections across the membrane. To our knowledge, this is the first time that signal transmission of abnormal activities (i.e., epileptiform discharge propagation) has been recapitulated in vitro.

**Figure 4.**
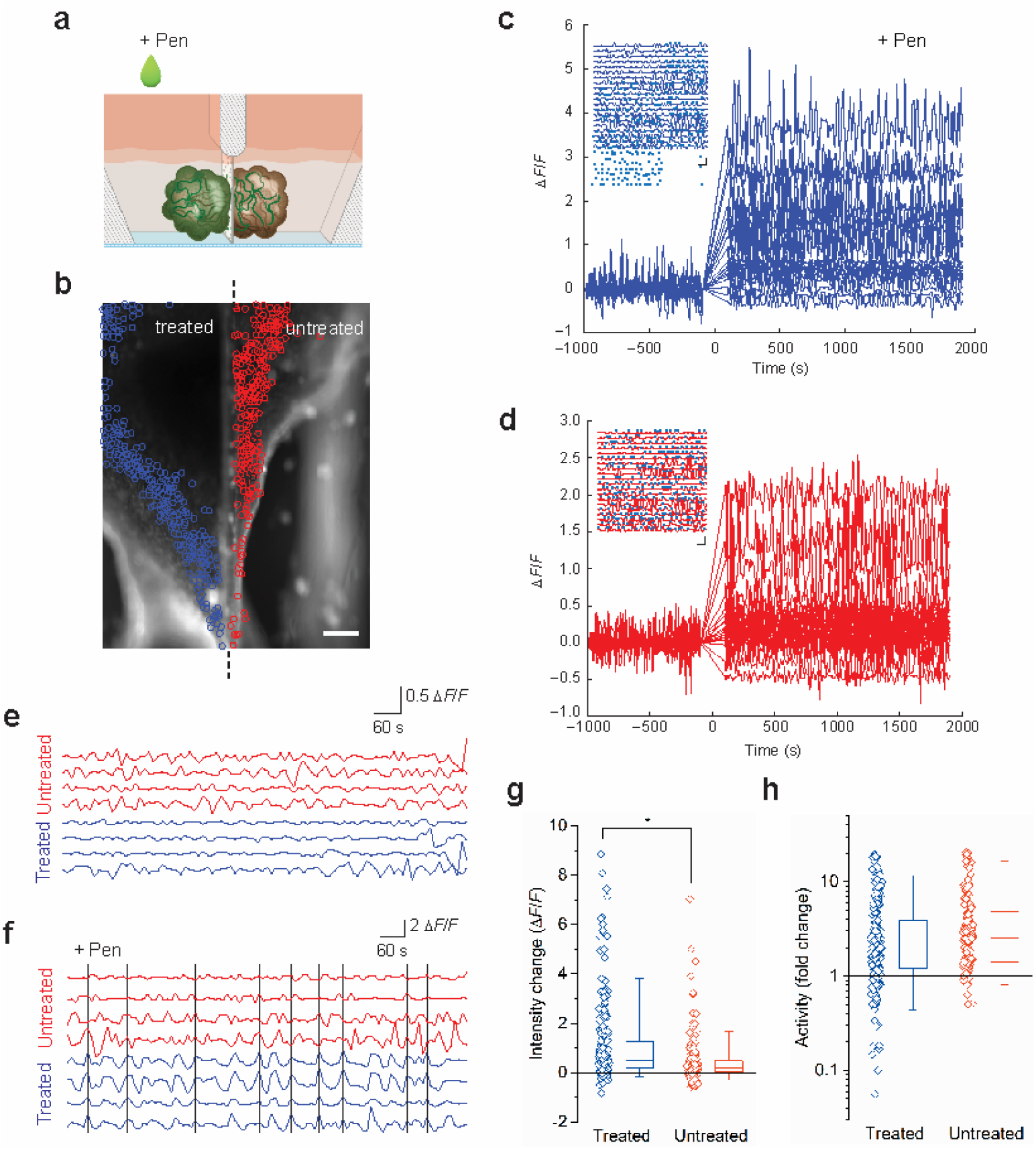
Discharge propagation between MARC-produced cerebral tissues. (**a,b**) Two cerebral tissues were separately formed in the two chambers of an iS3CC chip, separated by a membrane (black dashed line). One of the chambers (left, “treated”) was treated with Penicillin G (“Pen”), whereas the other (right, “untreated”) was not. The activity of 522 neurons were detected in the treated (blue circles) and untreated (red circles) tissues and analyzed by live calcium imaging (see also Supplementary Movie 3). Scale bar: 250 μm. (**c**,**d**) Time traces of normalized intensity (Δ*F/F*) from 20 representative neurons in the treated (**c**, blue) and untreated (**d**, red) are shown. Time 0 refers to the addition of Penicillin G (“Pen”). The inset show zoom-in views of the pre-treatment time traces. Data from the 20 cells were offset for clarity. Blue crosses indicate detected transient spikes. Vertical scale bars: 1 Δ*F*/*F*. Horizontal scale bars: 60 s. (**e**,**f**) Time traces of 4 cells in the treated (blue) and untreated (red) cerebral tissues pre- (**e**) and post-treatment (**f**) with Penicillin G. The black vertical lines indicate instances where all 4 cells in the treated tissue showed synchronized transient peaks. This synchronicity propagated ~45% of the time to the cells in the untreated tissue. (**g**,**h**) Quantification of the change in fluorescence intensity (**g**) and fold change in neuronal activity (**h**, log scale) induced by addition of Penicillin G in the treated (blue) and untreated (red) cerebral tissues. The symbols represent data for each cell, the boxes represent the median, 1^st^ and 3^rd^ quartiles, and the whiskers represent the 5^th^ and 95^th^ percentiles of the population data. Asterisk denotes statistically significant difference (Mann-Whitney U test, *p* < 10^−11^).

One of the key elements of the development of nervous systems is the formation of complex neuronal networks^27,28^. The engineered cerebral tissues in this study showed characteristics of mature neuronal networks, including synchronized influxes of extracellular calcium and modular functional connectivity patterns, demonstrating the formation of interconnected network within the intact tissues. Indeed, our culture method of promoting active reaggregation of cells and spheroids minimizes exogenous (mechanical) perturbations that may lead to cellular stress and unknown effects on tissue functionality ^29^. As such, the MARC-produced cerebral tissues can be used to study physiological and pathophysiological features of healthy and diseased neuronal networks. In general, neurological disorders are known to be accompanied by alterations in the network and functional interconnectivity between different brain regions^30^. Examples of these disorders include epilepsy, Alzheimer’s disease, schizophrenia, multiple sclerosis, depression, ASD, and traumatic brain injury. To further test the ability of our cerebral tissues to serve as a model of network interconnectivity disorders, we induced abnormal excessive discharges in the tissue cultured in one chamber of the iS3CC chip, which was found to lead to increased neuronal activity in the untreated tissue. These data indicate propagation of excessive discharges, which is the underlying mechanism of occurrence of an epileptic seizure and the target of most of current anti-epileptic drug treatments. Moreover, our study demonstrates a novel approach to develop and analyze interconnected brain tissue that opens a wide range of possibilities to mechanistically study clinically relevant interregional functional connectivity in lab grown 3D brain tissues.

## Acknowledgments

We thank Jan de Boer, Jaap den Toonder, Anton de Louw, and Remco van der Hofstad for insightful discussions, Jurgen Bulsink for help with the fabrication of the iS3CC chip molds, Serena Buscone for support with immunohistochemical staining, Jos Broers and Florence van Tienen (Maastricht University) for sharing the hiPS cells, and Koen Pieterse (ICMS Animation Studio) for help with the illustrations used in this manuscript. The authors acknowledge support from the Dutch Research Council (grant OCENW.XS2.017, to N.A.K.) and the Netherlands’s Ministry of Education, Culture, and Science (Gravitation program “Materials-Driven Regeneration”, grant 024.003.103, to N.A.K and C.V.C.B.).

## Author contributions

A.S., N.A.K., and C.V.C.B. developed the initial concept; A.S. developed the experimental approach and designed the culture devices, performed experiments, and analyzed data with input from N.A.K. and C.V.C.B.; A.P.A. conceived and supervised the network disorder study; N.A.K. developed and performed the network analysis; A.P.A., N.A.K., and C.V.C.B. provided support and supervised the project; A.S. and N.A.K. wrote the manuscript; All authors discussed results, contributed to revision of the manuscript, and approved the manuscript.

## Methods

### Cell culture

Human induced pluripotent stem cells (hiPSCs) were cultured in mTeSR medium (STEMCELL Technologies). The cell culture flasks were coated with Matrigel (hECS-qualified matrix, Corning, C354277) diluted in a 1:1 mixture of Dulbecco’s Modified Eagle’s Medium (DMEM, Gibco) and Ham’s F-12 Nutrient Mixture (Gibco) with a v/v ratio of 1:80 for 2 hours in an incubator at 37°C and 5% CO_2_. On the first day of culture, the hiPSCs were treated with 10 μM ROCK inhibitor Y-27632 (STEMCELL Technologies). The cells were washed using Dulbecco’s Phosphate-Buffered Saline (DPBS, Gibco) and the medium was refreshed daily until 80–100% confluence was reached.

### Neural induction

The cells were switched to neural induction medium containing 1:1 mixture of N2/B27 medium containing 10 ng/ml basic fibroblast growth factor (b-FGF), 1 μM Dorsomorphin dihydrochloride (Tocris, 3093), 10 μM SB431542 (Tocris, 1614), 100 ng/ml mouse recombinant Sonic Hedgehog (SHH)-C25II (Genscript, Z03050-50), and 10 μM CHIR99021 (Sigma, SML1046). N2 medium consisted of DMEM/F12 medium (Gibco) with 1× N2 supplement (Gibco, 17502048), 5 μg/ml insulin (Sigma, 19278), 1 mM L-Glutamine (Lonza, 17605E), 100 μM MEM-Non-Essential Amino Acid solution (NEAA) (Gibco), 100 μM 2-mercaptoathanol (Sigma, M3148), and 1:100 Penicillin-Streptomycin (Lonza, 17602E). B27 medium consisted of Neurobasal medium (Gibco) and 1× B27 supplement (Gibco, 17504044). The cells were washed daily using DPBS and maintained in induction medium.

### Supported reaggregation and tissue formation

#### Neural differentiation

After 1 week of neural induction, the cells were dissociated using Accutase (STEMCELL Technologies, 07920) and resuspended in growth factor reduced (GFR) Matrigel (Corning, 734-0269) and neural differentiation medium with a 70:30 v/v ratio at a density of 50,000 cells/chamber. Neural differentiation medium consisted of N2/B27 medium containing 10 ng/ml b-FGF, 20 ng/ml SHH-C25II, and 100 ng/ml human recombinant FGF8 (Gibco, PHG0184). The medium was refreshed daily for 7–10 days as the cells aggregated to form spheroids.

#### Pre-terminal differentiation and neural maintenance

Following spheroid formation, the medium was replaced with pre-terminal differentiation medium containing neural differentiation medium without b-FGF. After 10–15 days of culture, during which period the spheroids extended neurites and made extensive connections with each other, the medium was replaced with N2/B27 medium, entering the terminal differentiation phase. From this point, the medium was changed every other day.

### iS3CC device fabrication

The three-dimensional model of the device was built in Siemens NX (version NX10) software, from which the model of the negative mold was created. A polycarbonate (PC) negative mold was fabricated using micro-milling (Mikron wf 21C). PDMS silicon elastomer kit (Sylgard 184) was used to create the devices using soft lithography. A solution of silicon elastomer and curing agent with a weight ratio of 10:1 was mixed and degassed and then poured into the molds and cured in the oven at 80 °C for 3 hours. After that, polyethylene terephthalate (PET) porous membranes with a pore size of 8 μm (ThinCert, 657638) were cut into desired size and placed in the right position in the chip. Gluing and immobilization of the chip and the membrane on a 0.17 mm glass slide was applied using PDMS mixture with the same composition as mentioned earlier. The chips had a final cure in the oven at 110 °C for 2 hours.

### Cerebral tissue formation and maintenance in the iS3CC chips

The transparent PDMS devices were repeatedly washed using 70% ethanol followed by sterilization using UV light in the safety cabinets (3× 5 min). After neuronal induction phase, the cells were disassociated using Accutase and resuspended in 35 μl of a mixture of cold GFR-Matrigel and differentiation medium with a 70:30 v/v ratio at a final concentration of 50,000 cells per chamber. The chips containing cells and Matrigel-medium mixture were placed in a Petri dish to prevent contamination, since the chips have an open top, and kept in an incubator at 37°C and 5% CO_2_ for 5 min to polymerize the Matrigel mixture. After that, an additional 200 μl of differentiation medium was added to each chamber and the chips were placed back into the incubator. The medium was refreshed and the culture was continued as described above, with gentler handing to prevent damage to the gel and cells.

### Immunohistochemistry

The cerebral tissues were fixed in 3.7% paraformaldehyde for 2 hours at 4 °C and washed five times with PBS for 10 min. After that they were gently detached and taken out of the chambers and placed in cryomolds (Tissue-Tek). After that, OCT compound (Tissue-Tek) embedding medium was added to the cerebral tissues and snap-frozen on dry ice. Sections of 10 μm and 100 μm were created using cryotome (Microm, HM 550). For immunostaining, the sections were dried at room temperature and subsequently permeabilized for 10 min using 0.5% Triton X-100 in PBS and blocked 2×10 min using 10% normal donkey serum. After that, sections were incubated in the following primary antibodies diluted in PBS containing 1% normal donkey serum: rabbit polyclonal anti-Pax6 (1:200, Invitrogen, 42-6600), mouse monoclonal anti-beta-Tubulin-III, Tuj1 (1:200, Merck, MAB1637), chicken polyclonal to tyrosine hydroxylase, TH (1:200, Abcam, ab76442), rabbit monoclonal to Glutaminase (1:200, Thermofisher, 701965), mouse monoclonal anti vesicular glutamate transporter-I, VGLUT1 (1:200, Merck, AMAB91041), rat monoclonal anti-Dopamine Transporter, DAT (1:200, Abcam, ab5990), rabbit anti-VGAT (1:200, Merck, AB5062P), mouse anti-GFAP (1:200, Merck, G3893), chicken polyclonal anti-MAP2 (1:200, Abcam, ab75713), rabbit anti-OLIG2 (1:200, Merck, HPA003254). Fluorescence images of sections were obtained using a widefield epifluorescence microscope (Leica DMi8) equipped with a 10× / 0.32 HC PL Fluotar or a 20× / 0.4 HC PL Fluotar objective lens.

### Ca^2+^ imaging

Live calcium imaging was performed with the same widefield epifluorescence microscope, equipped with temperature, CO_2_, and humidity control. The tissues were incubated in the recommended concentrations of Fluo-4 direct according to the manufacturer (Molecular Probes, F10471) for 50 min in the incubator at 37 °C and 10 min at room temperature. After that, the tissues were washed five times with the neural maintenance medium. The calcium surges were recorded using an excitation of 488 nm and an emission of 530 nm every 10 seconds. Fluorescence images were obtained using a widefield epifluorescence microscope (Leica DMi8) equipped with either a 5× / 0.15 HC PL Fluotar or a 10× / 0.32 HC PL Fluotar objective lens.

### Penicillin treatment

After 15 minutes of calcium imaging in normal conditions, the chip was taken out of the live-imaging setup and 170 μl of the total volume of the treated chamber was replaced by a solution of Penicillin G sodium salt (Sigma, 13752) with a concentration of 100 mg/ml (equivalent to 2,8 × 10^4^ IU). The chip was quickly placed back in the same position and the live calcium imaging was continued. This process took approximately 3 min.

### Analysis of network interconnectivity

Neuronal activity was analyzed from the time-lapse images. To account for possible drifts or deformations of the tissues, the location of the cells was detected in each frame and linked across the frames using the Mosaic particle tracker plugin in ImageJ. The fluorescence intensity of each detected cell was calculated from the image intensity data using MATLAB. Occasional gaps in the time traces, due to the cells not being detected in certain frames, were filled using spline interpolation of the intensity values. The background fluorescence in each chamber was calculated in the same way using 10 arbitrarily selected cell-sized ROIs in the cell-free regions of each chamber, averaged, and subtracted from the real (cell) data. Further, the intensity values were corrected for imaging artifacts due to out-of-focus fluorescence by subtracting the mean intensity of an annular mask with an outer radius of 18 pixels and an inner radius of 9 pixels (i.e., size of the ROI) for each ROI.

Further analysis of the functional interconnectivity between neurons was performed using a custom-written script in MATLAB (version R2018b, The Mathworks Inc.). The normalized rate of change in fluorescence (ΔF/F) was calculated using (F_cell_ - F_min_)/F_min_ where F_cell_ is the mean fluorescence of a selected ROI measured in each frame and F_min_ the lowest measured mean fluorescence value of that ROI throughout the imaging window. For the analysis of the neuronal activity upon Penicillin treatment, ΔF/F was calculated as (F_cell_ - F_init_)/F_init_, where F_init_ is the fluorescence intensity of the ROI at the beginning of the live imaging, to capture the jump in fluorescence intensity due to addition of Penicillin. Transient spikes with minimum peak prominence larger than twice the magnitude of stochastic noise in the cell-free regions were detected using the ‘findpeaks’ function. To identify synchronized neuronal firings, the Pearson’s linear correlation coefficient *r* was computed for each pair of detected ROI. Following Eguíluz et al^11^, we defined two ROIs to be functionally connected when *r* > 0.6. To further assess the network modularity, we analyzed the functional connectivity using iterative Louvain community-detection algorithm^19,20^ with weighted edges to find the optimal network partitioning. Visualization of the network connectivity was realized using the graph plotting functions in MATLAB.

**Diffusion rate measurements in the iS3CC device** for analysis of relative diffusion rate, one of the chambers of the iS3CC device was loaded with 100 mg/ml of fluorescein sodium salt in Milli-Q water and the other chamber with pure Milli-Q. At different time points, samples of 5 μl were taken out from the pure Milli-Q chamber and added to a 96 well microplate to a final volume of 100 μl and fluorescent intensity was measured using a plate reader (Synergy HT). Based on calibration experiments, the final normalized concentrations in the MilliQ chamber indicated the relative diffusion rate over time.

## References

1. Quadrato, G. et al. Cell diversity and network dynamics in photosensitive human brain organoids. Nature 545, 48–53 (2017).

2. Giandomenico, S. L. et al. Cerebral organoids at the air–liquid interface generate diverse nerve tracts with functional output. Nat. Neurosci. 22, 669–679 (2019).

3. Birey, F. et al. Assembly of functionally integrated human forebrain spheroids. Nature 545, 54–59 (2017).

4. Paşca, A. M. et al. Functional cortical neurons and astrocytes from human pluripotent stem cells in 3D culture. Nat. Protoc. 12, 671–678 (2015).

5. Lancaster, M. A. et al. Guided self-organization and cortical plate formation in human brain organoids. Nat. Biotechnol. 35, 659–666 (2017).

6. Lancaster, M. A. & Knoblich, J. A. Organogenesis in a dish: Modeling development and disease using organoid technologies. Science (80-.). 345, 1247125. (2014).

7. Eiraku, M. et al. Article Self-Organized Formation of Polarized Cortical Tissues from ESCs and Its Active Manipulation by Extrinsic Signals. Stem Cell 3, 519–532 (2008).

8. Valdeolmillos, M. & Moya, F. Leading Process Dynamics During Neuronal Migration. in Cellular Migration and Formation of Neuronal Connections (eds. Rubenstein, J. L. R. & Pasko, R.) 245–260 (Academic Press, 2013). doi:10.1016/B978-0-12-397266-8.00025-9.

9. Ohtaka-Maruyama, C. et al. Synaptic transmission from subplate neurons controls radial migration of neocortical neurons. Science (80-.). 360, 313–317 (2018).

10. Robinson, H. P. et al. Periodic synchronized bursting and intracellular calcium transients elicited by low magnesium in cultured cortical neurons. J. Neurophysiol. 70, 1606–1616 (1993).

11. Eguiluz, V. M., Chialvo, D. R., Cecchi, G. A., Baliki, M. & Apkarian, A. V. Scale-Free Brain Functional Networks. Phys. Rev. Lett. 94, 018102 (2005).

12. Feldt, S., Bonifazi, P. & Cossart, R. Dissecting functional connectivity of neuronal microcircuits: experimental and theoretical insights. Trends Neurosci. 34, 225–236 (2011).

13. Batista, C. A. S., Batista, A. M., Pontes, J. A. C. De, Viana, R. L. & Lopes, S. R. Chaotic phase synchronization in scale-free networks of bursting neurons. Phys. Rev. E 76, 016218 (2007).

14. Grinstein, G. & Linsker, R. Synchronous neural activity in scale-free network models versus random network models. Proc. Natl. Acad. Sci. 102, 9948–9953 (2005).

15. Barabási, A. & Reka, A. Emergence of Scaling in Random Networks. Science (80-.). 286, 509–512 (1999).

16. He, B. J. Scale-free brain activity: past, present and future. Trends Cogn Sci 18, 480–487 (2015).

17. Mcintosh, A. R. Towards a network theory of cognition. Neural Networks 13, 861–870 (2000).

18. Cohen, J. R. & Esposito, M. D. The Segregation and Integration of Distinct Brain Networks and Their Relationship to Cognition. J. Neurosci. 36, 12083–12094 (2016).

19. Blondel, V. D., Guillaume, J., Lambiotte, R. & Lefebvre, E. Fast unfolding of communities in large networks. J. Stat. Mech. theory Exp. 10, P10008 (2008).

20. Pedersen, M., Zalesky, A., Omidvarnia, A. & Jackson, G. D. Multilayer network switching rate predicts brain performance. Proc. Natl. Acad. Sci. 115, 13376–13381 (2018).

21. Heuvel, M. P. Van Den, Stam, C. J., Kahn, S. & Pol, H. E. H. Efficiency of Functional Brain Networks and Intellectual Performance. J. Neurosci. 29, 7619–7624 (2009).

22. Fruchterman, T. M. J. & Reingold, E. M. Graph Drawing by Force-directed Placement. Software-practice Exp. 21, 1129–1164 (1991).

23. Johnson, H. C. & Walker, E. Intraventricular penicillin: a note of warning. J. Am. Med. Assoc. 127, 217–219 (1945).

24. Avoli, M. & Jefferys, J. G. R. Models of drug-induced epileptiform synchronization in vitro. J. Neurosci. Methods 260, 26–32 (2016).

25. Rossokhin, A. V., Sharonova, I. N., Bukanova, J. V., Kolbaev, S. N. & Skrebitsky, V. G. Block of GABAA receptor ion channel by penicillin: Electrophysiological and modeling insights toward the mechanism. Mol. Cell. Neurosci. 63, 72–82 (2014).

26. Treiman, D. M. GABAergic mechanisms in epilepsy. Epilepsia 42, 8–12 (2001).

27. Katz, L. C. & Shatz, C. J. Synaptic Activity and the Construction of Cortical Circuits. Science (80-.). 274, 1133–1138 (1996).

28. Zhang, L. I. & Poo, M. M. Electrical activity of neural circuits. Nat. Neurosci. 4, 1207–1214 (2001).

29. Bhaduri, A. et al. Cell stress in cortical organoids impairs molecular subtype specification. Nature 578, 142–148 (2020).

30. Lubrini, G., Martín-Montes, A., Díez-Ascaso, O. & Díez-Tejedor, E. Brain disease, connectivity, plasticity and cognitive therapy: A neurological view of mental disorders. Neurologia 33, 187–191 (2018).

