## Supplementary material for "Human cerebral tissues created via active cellular reaggregation produce functionally interconnected 3D neuronal network to mimic pathological circuit disturbance"

### Supplementary Figures

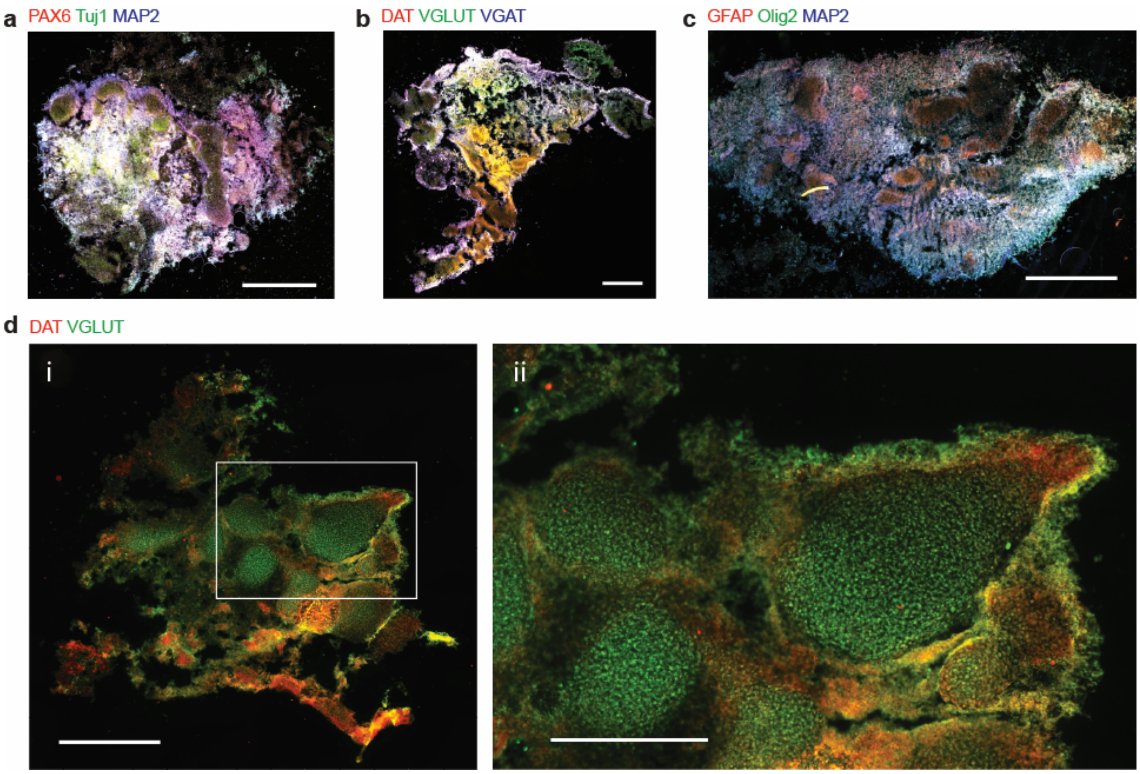


**Figure S1** Immunohistochemical co-staining of multiple markers on 100 μm sections of MARC-produced cerebral tissues at Day 90 revealed the overall distribution and regionalization of distinct neuronal cell types. Co-staining of neural progenitor cells (NPCs; PAX6 in red), early neurons (Tuj1 in green) and mature neurons (MAP2 in blue) (**a**), co-staining of dopamine transporter (DAT in red) vesicular transporters of glutamine (VGLUT in green) and GABA (VGAT in blue) (**b**) co-staining GFAP (red), Olig2 (green) and MAP2 (blue) (**c**) and co-staining of dopamine transporter (DAT in red) and vesicular glutamine transporter (VGLUT in green) in particular (**d i and ii)** indicate the localized co-expression of these markers suggesting regionalization of these cell types in the cerebral tissues. Scale bars: 1 mm.

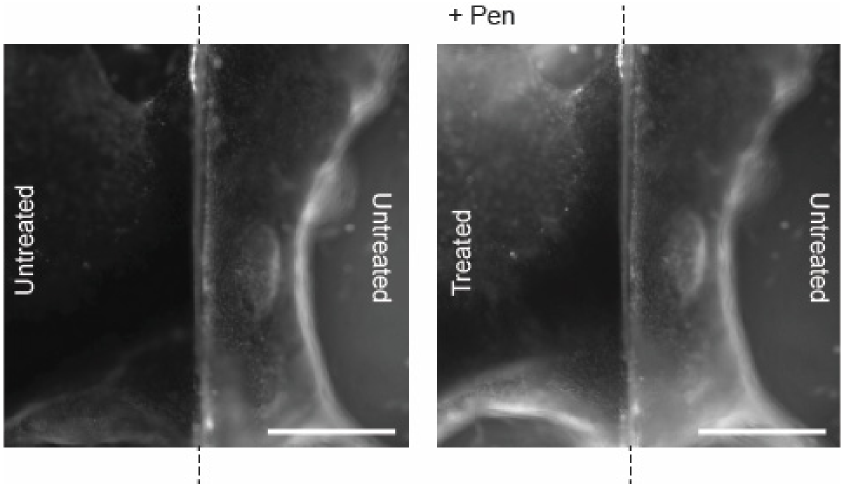


**Figure S2** Immediate fluorescence increase in the treated chamber upon Penicillin G treatment. Two cerebral tissues were separately formed via MARC protocol in the two chambers of an iS3CC chip, separated by a membrane (black dashed line). One of the chambers (left, “treated”) was treated with Penicillin G (“Pen”), whereas the other (right, “untreated”) was not treated. Scale bar: 1mm.

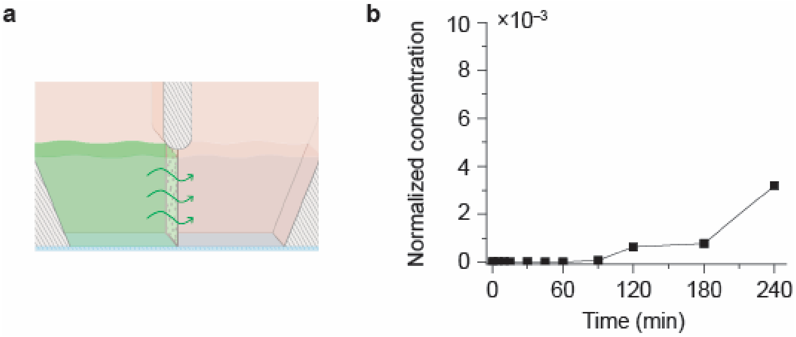


**Figure S3** Measurement of particle transfer between the chambers of the iS3CC chip across the porous membrane. Fluorescein sodium salt with comparable molecular weight (376.27 g/mol) to Penicillin G sodium salt (367.37 g/mol), was added to one of the chambers of the iS3CC with the exact final concentration as Penicillin treatment (100 mg/ml) and the fluorescence of water in the other chamber was measured overtime using a plate reader (see Methods).

### Caption to Supplementary Movies

**Movie 1**

Time lapse recording of live fluorescent calcium imaging of a MARC-produced cerebral tissue (week 4).

**Movie 2**

Time lapse recording of of live fluorescent calcium imaging of neurite connections between two cerebral tissues across the porous membrane of the iS3CC (week 6).

**Movie 3**

Time lapse recording of live fluorescent calcium imaging of two interconnected cerebral tissues in the iS3CC before and after Penicillin treatment in one of the chambers (week 6).
